# A multifunctional R package for identification of tumor-specific neoantigens

**DOI:** 10.1101/869388

**Authors:** Takanori Hasegawa, Shuto Hayashi, Eigo Shimizu, Shinichi Mizuno, Atsushi Niida, Rui Yamaguchi, Satoru Miyano, Hidewaki Nakagawa, Seiya Imoto

## Abstract

It is known that some mutated peptides, such as those resulting from missense mutations and frameshift insertions, can bind to the major histocompatibility complex and be presented to antitumor T-cells on the surface of a tumor cell. These peptides are termed neoantigen and it is important to understand this process for cancer immunotherapy. Here, we introduce an R package that can predict a list of potential neoantigens from a variety of mutations, which include not only somatic point mutations but insertions, deletions, and structural variants. Beyond the existing applications, this package is capable of attaching and reflecting several additional information, *e.g*., wild-type binding capability, allele specific RNA expression levels, single nucleotide polymorphism information, and combinations of mutations to filter out infeasible peptides as neoantigen.

**Availability:** The R package is available at http://github/hase62/Neoantimon.

## Introduction

Recent technological advances in massively parallel sequencing have enabled identification of genetic variants, *e.g.*, single nucleotide variants (SNVs) and insertions or deletions (Indels), in individual cancer patients. Furthermore, substantial evidence indicates that tumor-specific peptides that result from such variations can bind to a major histocompatibility complex (MHC) molecule and be presented to antitumor T-cells on the surface of a tumor cell. Identification of such tumor-specific peptides, termed neoantigens, has been receiving increasing attention because of its numerous potential applications in cancer immunotherapy.

To identify the possible presence of neoantigens in individual tumor, we have to predict whether mutated peptides can bind to the patient’s human leukocyte antigens (HLAs). Several computational methods such as netMHCpan [4] have been proposed to predict the binding capability, including binding affinity and percentage rank of affinity. To apply these methodologies, we need to determine the patient’s HLA types and prepare a list of tumor-specific peptides obtained from sequencing data. Designing such peptides requires not only mutation information, but also the reference sequences with their coding protein information because they are fractions of expressed mutated proteins. Even after the prediction of the binding capability of such mutated peptides, further classification filters should be applied to the selection, *e.g*., a comparison in binding affinity between wild-type and mutated peptides and the evaluation of allele specific RNA expression levels.

To automate this process and easily identify tumor-specific neoantigens, some computational tools have been developed [2,3]. These tools greatly help to obtain predicted results; however, a few tools can employ mutation data (such as variant call format (vcf) files) in a local analytical environment such as the R. In addition, the existing tools are applicable only for the prediction of HLA Class I binding and for handling of SNVs and Indels at best, but not for the prediction of HLA Class II binding and handling of structural variants (SVs). Moreover, they lack detailed considerations, *e.g*., reflecting SNVs on the frameshift regions and single nucleotide polymorphisms (SNPs) to generated peptides. To address these requirements, we developed an easy and multifunctional R package that can produce a list of candidate neoantigens (for HLA Class I and II) caused by SNVs, Indels, and SVs. It can automatically construct mutated peptides from vcf files or mutant RNA sequences and calculate their binding capability to the corresponding HLAs with some information for filtering. This tool has been used in the Mitochondrial Genome and Immunogenomics Working Group in the PanCancer Analysis of Whole Genomes (PCAWG) project [5].

## Materials and Methods

### Input files

This package requires the two following inputs: (i) an annotated vcf file generated using, *e.g.*, ANNOVAR [7] and (ii) a list of HLA types. Otherwise, (i) can be replaced by either non-annotated vcf file with annotating option or (iii) a list of mutant RNA sequences in association with the corresponding gene symbol or NM IDs to filter out wild-type peptides. Users can optionally provide SNPs data to reflect them to mutated peptides, and also RNA expression profiles with RNA bam files and copy number variation data to attach bulk and allele specific RNA expression levels and tumor sub-clonality for filtering. Note that vcf files must conform to the BND format for the evaluation of potential fusion transcripts on the basis of SVs. The comparison of the functions among pVACseq [3] and MuPeXI [2], and our R package (neoantimon) is displayed in Table 1. One can install this package from a GitHub repository and use it in the R environment on Mac/Linux.

**Table 1.**
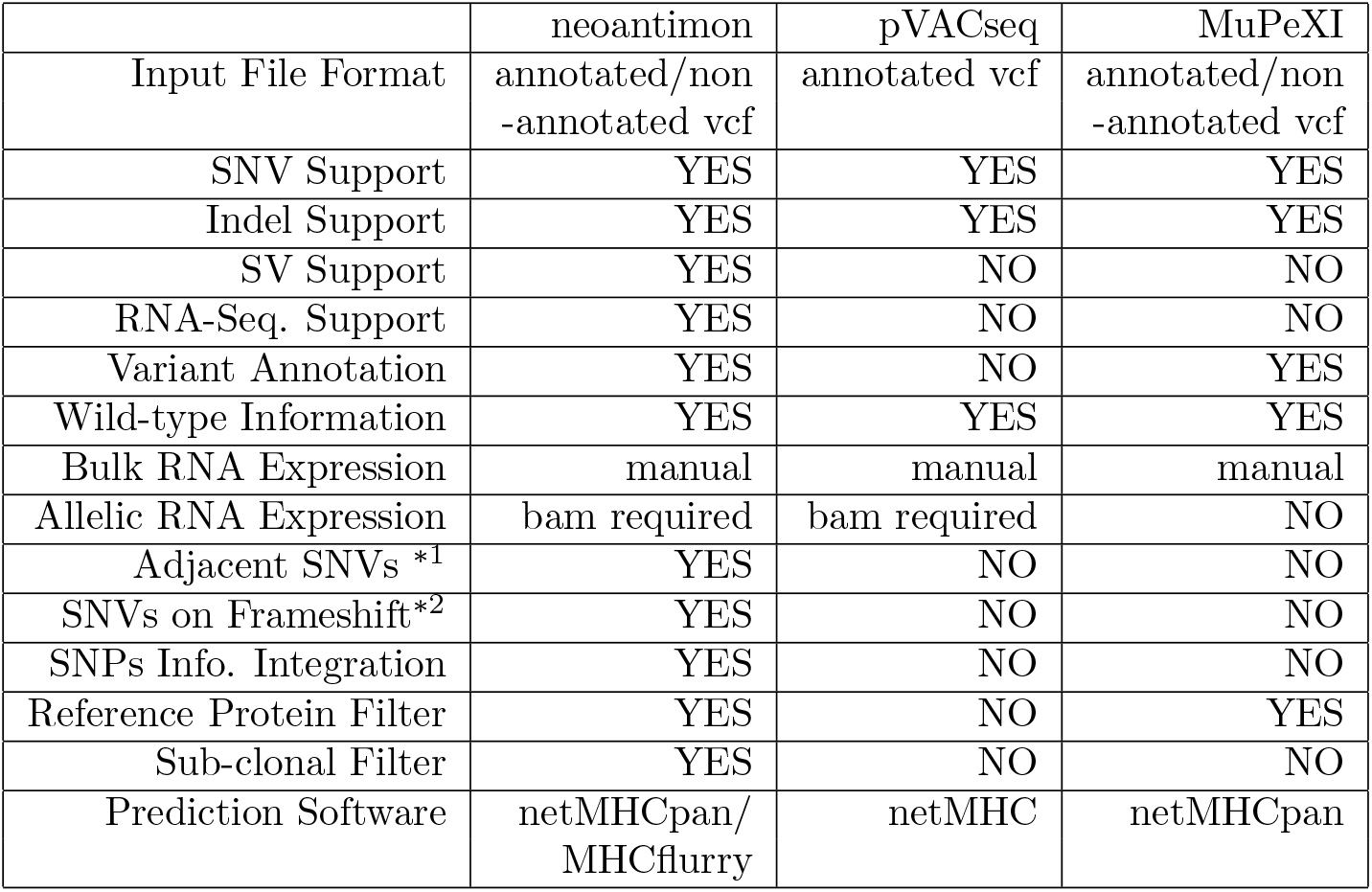
A comparison table of the functions among pVACseq, MuPeXI, and Neoantimon. ‘‘manual” means that users manually upload RNA expression files. ^*1^ and ^*2^ are the considerations of cases where SNVs are occurring next to each other and among frameshift regions generated by Indels, respectively.

### Output files

This package generates FASTA files consisting of mutated and corresponding wild-type (for SNVs) peptides according to the RefSeq trasncript sequences, and an integrated output file including peptide–MHC binding capability estimated by NetMHCpan4.0 [4] or MHCflurry [6], and NetMHCIIpan3.2 [1]. In the application to frameshift Indels and potential fusion transcripts, all mutated peptides generated from the mutation position to the stop codon are constructed. The integrated output file includes (1) the HLA type, (2) mutation position, (3) gene symbol and NM_ID, (4) exon start and end positions, (5) amino acid changes, (6) total read depth and variant allele frequency, (7) wild-type (if exists) and mutated peptide sequences, (8) their IC50 and percentages of rank affinity, (9) corresponding bulk RNA expression and variant allele frequency at the mutation position, (10) copy number of alleles A and B, (11) tumor sub-clonality as cancer cell fraction probability, and (11) additional information, *i.e*., flags for indicating the application of adjacent SNVs, SNVs in the frameshift region, and SNPs. Simple pictures of input and output files of this package are illustrated in Fig. 1.

**Figure 1:**
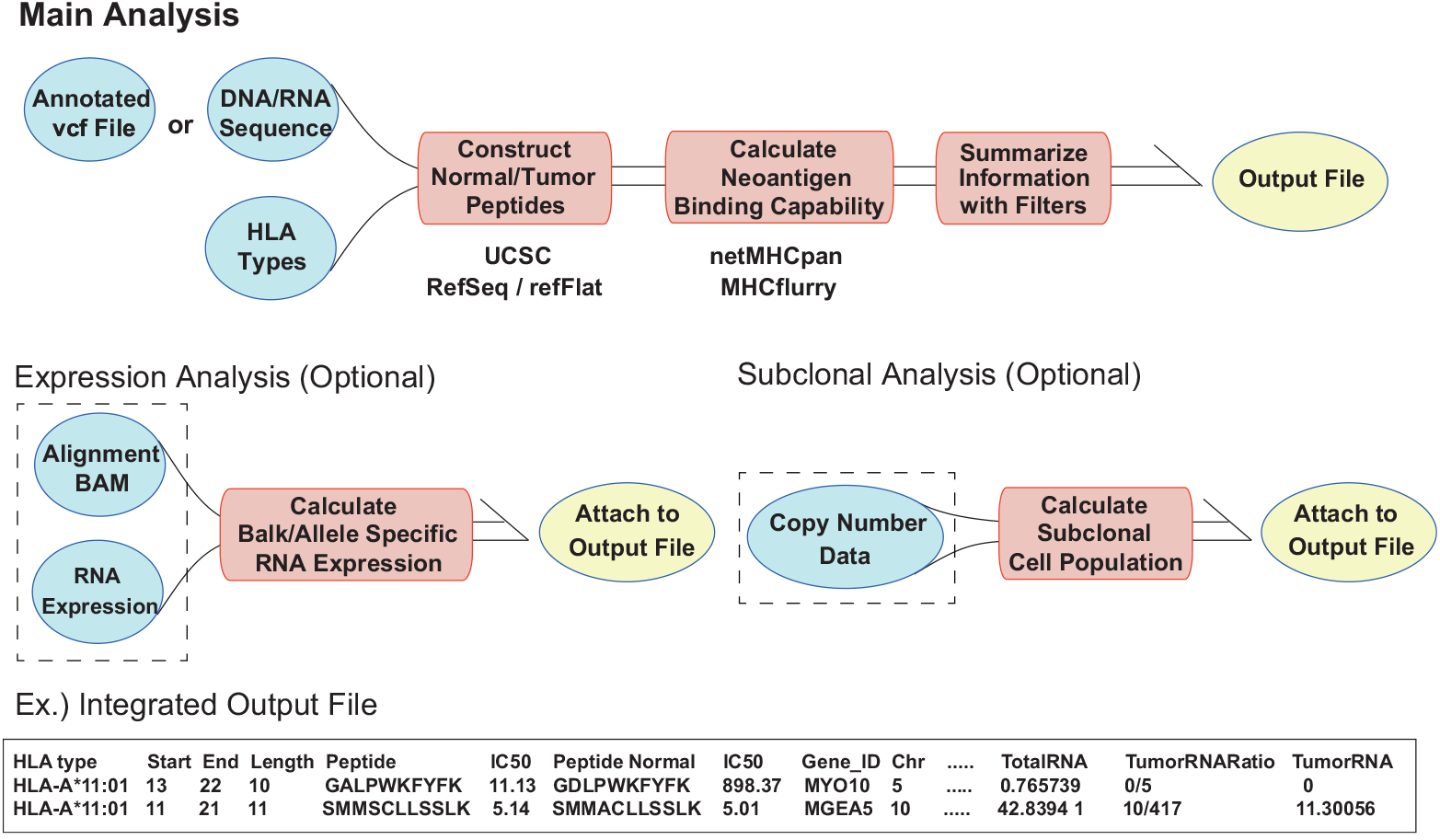
An overview of the input and output file information of the package. Green circles with and without dotted rectangles are optional and required inputs, respectively. Red rectangles and yellow circles are intermediate processes and the output files, respectively.

## Conclusions

We developed an R package generating candidate neoantigens from variety of mutations, *i.e*., SNVs, Indels, and SVs, and mutant RNA sequences. Beyond previously developed platforms, it can cover specific cases and include additional information for filtering. The package, documentation, and sample analysis are available at http://github/hase62/Neoantimon, and an analysis result in PCAWG project is also available [5].

